# *In Silico* Analysis of *B3GALTL* Gene Reveling 13 Novel Mutations Associated with Peters’-plus syndrome

**DOI:** 10.1101/2020.03.21.000695

**Authors:** Abdelrahman H. Abdelmoneim, Arwa A. Satti, Miysaa I. Abdelmageed, Naseem S. Murshed, Nafisa M. Elfadol, Mujahed I. Mustafa, Abdelrafie M. Makhawi

**Author notes:** Corresponding author: Abdelrahman H. Abdelmoneim.

## Abstract

**Background:** Peters’-plus syndrome is a rare autosomal recessive disorder, which is characterized by a specific malformation of the eye that includes corneal opaqueness and iridocorneal adhesions (Peters’ anomaly) along with other systemic manifestations. Furthermore, various researches report the association between B3GALTL gene and Peters’-plus syndrome. In the current work we aim to analyze the deleterious SNPs in B3GALTL gene that predispose to Peters’-plus syndrome.

**Method:** the associated SNPs of the coding region of the B3GALTL gene was acquired from National Center for Biotechnology Information and then analyzed by eight softwares (SIFT, Polyphen2, Proven, SNAP2, SNP@GO, PMut, Imutant and Mupro). The physiochemical properties of the resulted SNPs were then analyzed by Hope project website and visualized by chimera software.

**Result:** Thirteen novel mutations (Y172C, A222V, C260R, C260Y, D349G, I354K, R377C, G379C, G393R, G393E, G395E, G425E, R445W) are discovered in B3GALTL gene to cause deleterious effects leading to the development of Peters’-plus syndrome.

**Conclusion:** Thirteen novel mutations in B3GALTL gene are predicted to cause Peters’-plus syndrome.

## 1. Introduction

Peters Plus syndrome (MIM 261540) is a rare disorder with autosomal-recessive inheritance involving systemic and ocular abnormalities characterized by short stature, Genitourinary tract and anterior chamber abnormalities, and distinctive facial features like cleft lip and palate.(1-3) Of the abnormality reported in Peters plus syndrome, Peters anomaly is regarded as the most common ocular disorder which involves the thinning and clouding of the cornea and iridocorneal adhesions causing blurred vision. Other eye abnormalities, such as glaucoma and cataracts are also common.(4, 5) The diagnosis of Peters plus syndrome is a clinical diagnosis that can be confirmed by identification of B3GLCTgene mutations on molecular genetic testing. Treatment is mainly surgical with symptomatic treatment of complication.(6)

Peters plus syndrome is caused by mutations in the B3GLCT gene found in13q12.3 region, which encode for beta 3-glucosyltransferase enzyme that catalyzes glycosylation reactions through beta1-3 glycosidic linkage to O-linked fucose on thrombospondin type 1 repeats (TSRs).(7-9) TSRs are found in extracellular proteins and they plays important role in development, including angiogenesis and neurogenesis.(10) Mutations in the B3GLCT gene results in a short and nonfunctional enzyme that lacks this catalytic activities. Until now the molecular processes that leads to the signs and symptoms of the syndrome is still not well undrestood,(11) but impaired glycosylation likely disrupts the function of the proteins, which may contribute to the variety of features.(12) Absent mutation in B3GLCT gene in the present of other typical symptoms and sign are termed Peters plus-like syndrome.(13, 14)

Bioinformatics has become an extremely cost effect method for analyzing the vast amount of genetic data discovered in the recent years.(15) Better data sharing and analysis software are underdeveloped which is expected to yield more accurate and trustworthy results that will eventually transforms the landscape of healthcare.(16) Single-nucleotide polymorphisms (SNPs) are the most common form of human genetic variation.(17) Most single nucleotide polymorphisms (SNPS) that occur on genes are non-deleterious thus identify the disease causing SNPs is a complicated task,(18) and this highlights the importance behind the use of bioinformatics tools to discriminate between neutral and disease causing variant.

Most studies that is performed in our target focused in the conformation of the association of the gene to the disease (B3GLCT to peters plus syndrome).(19, 20) while other studies reported different type of mutation in B3GLCT gene including frameshift, nonsense, missense and splicing mutations with c.660+1G>A donor splice site mutation recognized as the most common mutation identified.(2, 21, 22) However our study focused on the determination of the most deleterious nonsense single nucleotide polymorphism of B3GLCT gene. This study will help in understanding the disease etiology, genotype to phenotype correlation and carrier detection since the disease is recessive disorder,(23, 24) in which affected people inherit one mutant copy from each parent. The value of determining the most deleterious SNPs mutation lies in the fact that carriers of autosomal recessive condition don’t show signs or symptoms (silent carriers) we believe our study will have a huge diagnostic values especially in families with disease history.

## 2. Material and method

Using dbSNP from the National Center of Biotechnology Information (NCBI), SNPs were retrieved, the interactions of B3GLCT with other genes were investigated using GeneMANIA, then functional analysis was performed using different online insilico analysis software as SIFT, PolyPhen-2, PROVEN, SNPs & GO and PMut. Stability analysis was performed using I_Mutant 3.0 and Mupro respectively, after that; the effect of SNPs on the structure was predicted using Project HOPE. Finally, the 3D structure of the protein was modeled using RaptorX and the altered amino acids were visualized using UCSF Chimera.

### 2.1. Data mining

SNPs were retrieved using dbSNP from National Center of Biotechnology Information(NCBI) (http://www.ncbi.nlm.nih.gov/SNP/). In addition, FASTA format of the reference sequence of human protein was collected from UnipProt database.

### 2.2. SIFT Server

It is an online in silico analysis tool which predicts the possible impact of amino acid substitutions on protein function using sequence homology. It gives scores for each amino acid from 0 to 1.SIFT scores < 0.05 are expected to be damaging, otherwise it’s considered to be tolerant. The software interpret the homologous sequence using SWISS_Prot and TrEMBL. (Available at: http://sift.bii.a-star.edu.sg/). ^(25-28)^

### 2.3. Provean Server

Its trained functional analysis tool which calculate whether amino acid substitution affects protein function or not.

The variant is expected to have a deleterious effect if PROVEN score < −2.5. If the score is higher than 2.5 the variant is expected to have a neutral effect (Available at: http://provean.jcvi.org/index.php). ^(29, 30)^

### 2.4. Polyphen-2 Server

It’s an online analysis software which predict the effect of an amino acid substitution on the structure and function of proteins using multiple sequence alignment, it estimates the position specific independent count score (PSIC) for every variant. The higher the PSIC value the higher the impact of the amino acid on the protein.

Prediction results classified as probably damaging (values are more frequently 1), possibly damaging or benign (values range from 0 to 0.95)(Available at: http://genetics.bwh.harvard.edu/pph2/). ^(31-33)^

### 2.5. SNAP2

It’s an online in silico functional analysis tool which differentiate between neutral and effect SNPs. Considered as one of the most reliable methods with 83% accuracy. The most important input signal for the prediction is the evolutionary information taken from an automatically generated multiple sequence alignment. Also structural features such as predicted secondary structure and solvent accessibility are considered. Although it take longer time than the previous software to get the result but it has a significant accuracy level (a sustained two-state accuracy (effect/neutral) of 82%).(34) It is available at https://rostlab.org/services/snap2web/.

### 2.6. SNPs & Go server

It’s an online in silico functional analysis software which predict whether a variation is disease related or not .it collects information from function as encoded in Gene Ontology term, evolutionary information. It under perform other available predictive methods as PhD-SNP and Panther. (if disease probability >0.5 mutation is considered as disease causing nsSNP). (Available at: http://snps-and-go.biocomp.unibo. it/snps-and-go/). ^(35)^

### 2.7. PMUT Server

It’s an online functional analysis tool for prediction of the effect of SNP alteration on proteins. With 80% accuracy in calculation of the compulsive feature of each SNP grounded on the practice of neural networks. It is available at (http://mmb.irbbarcelona.org/PMut). ^(36)^

### 2.8. Protein Stability analysis

#### I-Mutant

A online web-tool that is used for the prediction of protein stability changes upon single point mutations, determining whether the mutation increases or decreases the protein stability. After insertion of reference sequencing and the required position and related mutant amino-acid, a result page will appears with the desired prediction. Acquiring result may take some time depending on the server load. (Available at: http://gpcr.biocomp.unibo.it/cgi/predictors/I-Mutant2.0/I-Mutant2.0.cgi). ^(37)^

#### MUpro

It’s a collection of machine learning programs that predict how single amino acid substitutions affect protein stability. It was developed based on two machine learning methods: Neural Network and Support Vector Machine. Outcomes is either decreased stability if MUpro score <0 or increased stability if the score >0.(38)

### 2.9. GeneMANIA

It’s a web interface which predicts the relationship between related sets of genes together using associate data. Associate data includes genetic interaction and protein, co-expression, pathway, protein domain similarity. (Available at: (http://www.genemania.org/). ^(39)^

### 2.10. Modeling nsSNP locations on protein structure

#### Hope project

An Online software that analyze the structural and functional effects of point mutation. Project Hope collect information from a wide range of information including calculation on the 3d coordinates of protein by WHAT IF web server, prediction by DA services and sequence annotation from UniProt database.

Outcomes are presented in the form of a report with figures, test and annotation (40). It is available at http://www.cmbi.ru.nl/hope/.

#### Chimera

It is a visualization analysis tool of 3D structures, docking analysis and other analysis tools. It’s used to visualize the predicted 3D structure which was modeled by RaptorX, and compare the amino acid alteration of the original amino acid with the mutated one to see the impact that can be produced. Availabe at: http://www.cgl.ucsf.edu/chimera). ^(41)^

## 3. Results

A total 371 missense mutation were retrieved from dbSNP from the NCBI database, then SNPs were submitted to SIFT, PROVEN, PolyPhen-2 and SNAP2 respectively. SIFT predicted 262 deleterious SNPs, PROVEN predicted 172 deleterious SNPs, PolyPhen-2 predicted 172 probably damaging SNPs and 50 possibly damaging SNPS and finally SNAP2 predicted 167 deleterious SNPs.

Then SNPs were filtered based on the 4 previous software and the number decreased rapidly into 13 SNPs which has positive results in all of them. Then further investigation were carried out on these SNPs and their effect on the functional levels by SNAP & GO,PhD-SNAP and PMut as shown in table 2. Finally we run stability analysis using I-mutant 3.0 and MUpro respectively, 7 SNPs were found to decrease the stability by I-mutant 3.0 while all the 13 SNPs decrease the stability according to MUpro.

**Table 1:**
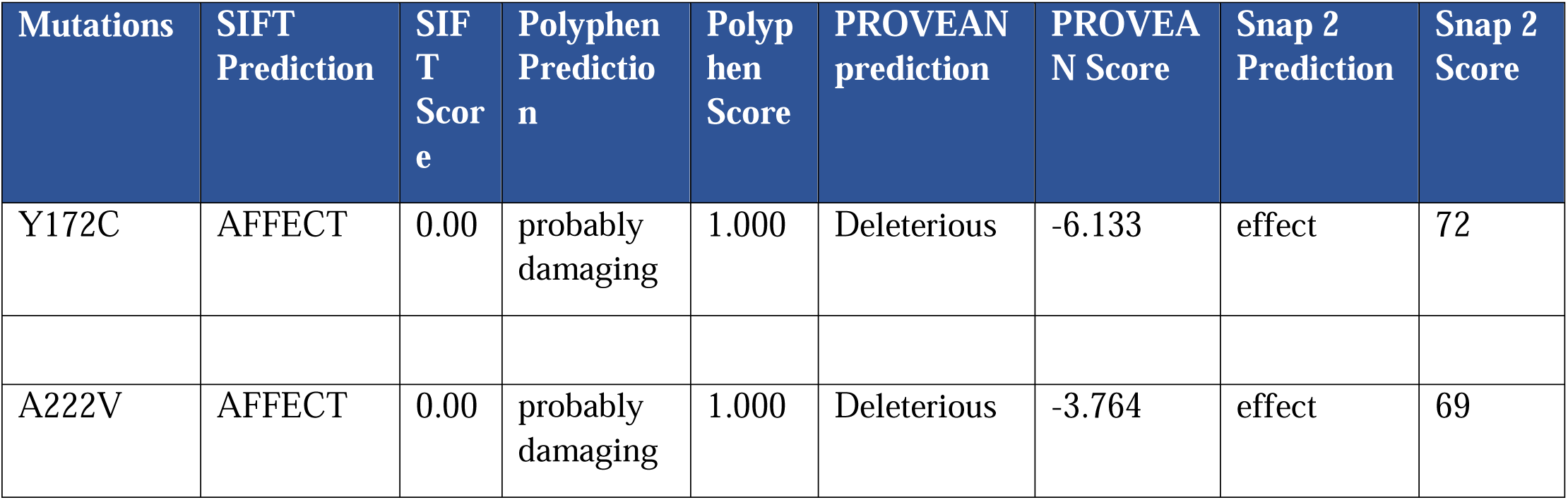
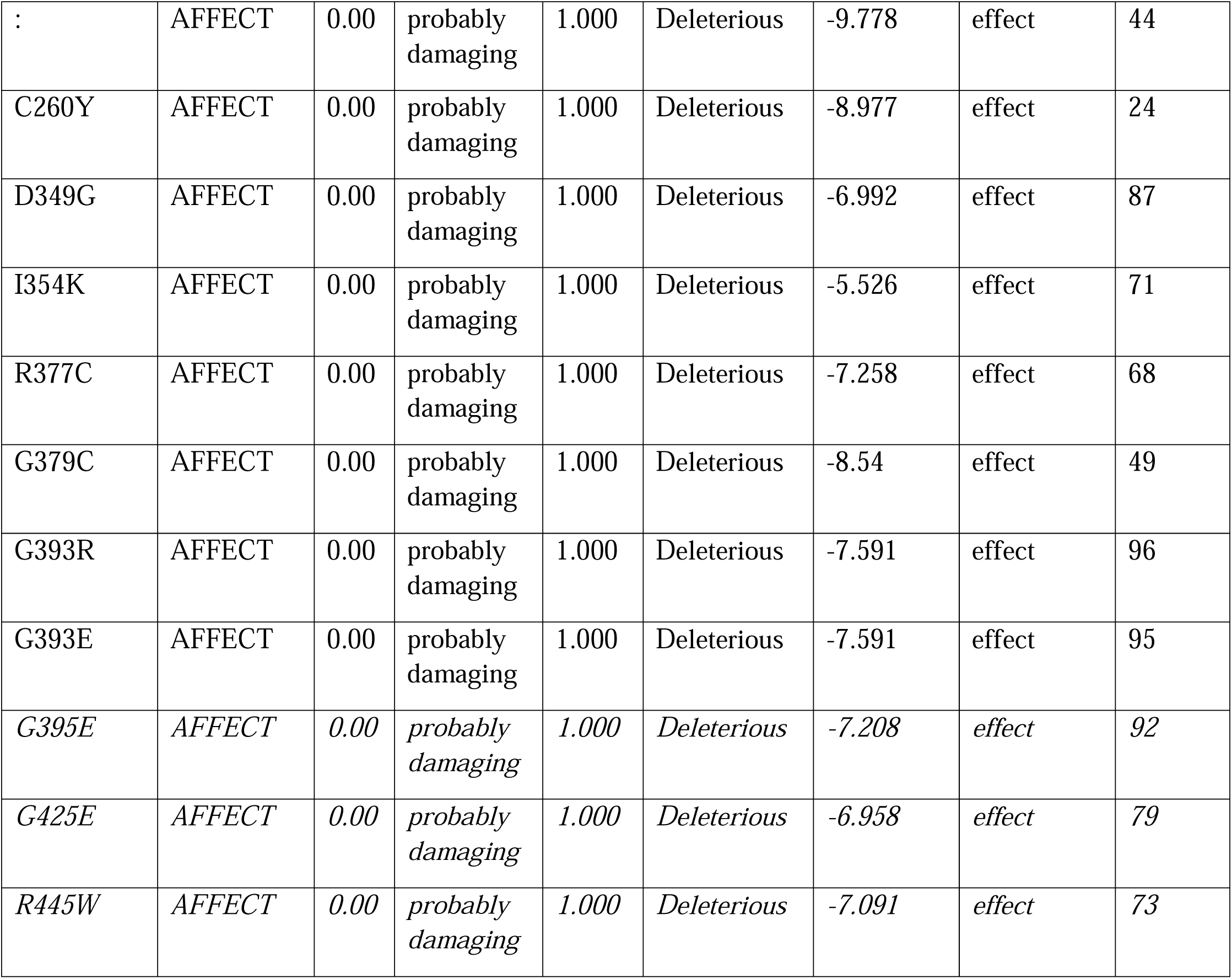
Affect or damaging nsSNPs predicted by various software.

**Table 2:** Pathogenic nsSNPs predicted by various software.

| Mutations | SNPs&GO Prediction | RI | Probability | PHD Prediction | RI | Probability | P-mut Prediction | Probability |
| --- | --- | --- | --- | --- | --- | --- | --- | --- |
| Y172C | Disease | 0 | 0.522 | Disease | 3 | 0.659 | Disease | 0.55<br>(81%) |
| A222V | Disease | 1 | 0.558 | Disease | 7 | 0.829 | Disease | 0.56<br>(81%) |
| C260R | Disease | 3 | 0.649 | Disease | 5 | 0.733 | Disease | 0.81 |
|  |  |  |  |  |  |  |  | (89%) |
| C260Y | Disease | 2 | 0.584 | Disease | 5 | 0.733 | Disease | 0.75<br>(87%) |
| D349G | Disease | 6 | 0.791 | Disease | 8 | 0.893 | Disease | 0.85<br>(91%) |
| I354K | Disease | 4 | 0.677 | Disease | 2 | 0.624 | Disease | 0.85<br>(91%) |
| R377C | Disease | 5 | 0.749 | Disease | 6 | 0.797 | Disease | 0.65<br>(84%) |
| G379C | Disease | 5 | 0.76 | Disease | 6 | 0.791 | Disease | 0.78<br>(88%) |
| G393R | Disease | 7 | 0.832 | Disease | 8 | 0.892 | Disease | 0.87<br>(91%) |
| G393E | Disease | 6 | 0.824 | Disease | 8 | 0.901 | Disease | 0.83<br>(90%) |
| G395E | Disease | 8 | 0.878 | Disease | 8 | 0.916 | Disease | 0.87<br>(92%) |
| G425E | Disease | 2 | 0.58 | Disease | 7 | 0.840 | Disease | 0.82<br>(90%) |
| R445W | Disease | 3 | 0.633 | Disease | 5 | 0.766 | Disease | 0.70<br>(86%) |

In this study B3GLCT gene was found to have an association with 20 other different genes using geneMAINA software. Co expression and physical interaction with other related genes is shown in Figure (2)

**Figure (1):**
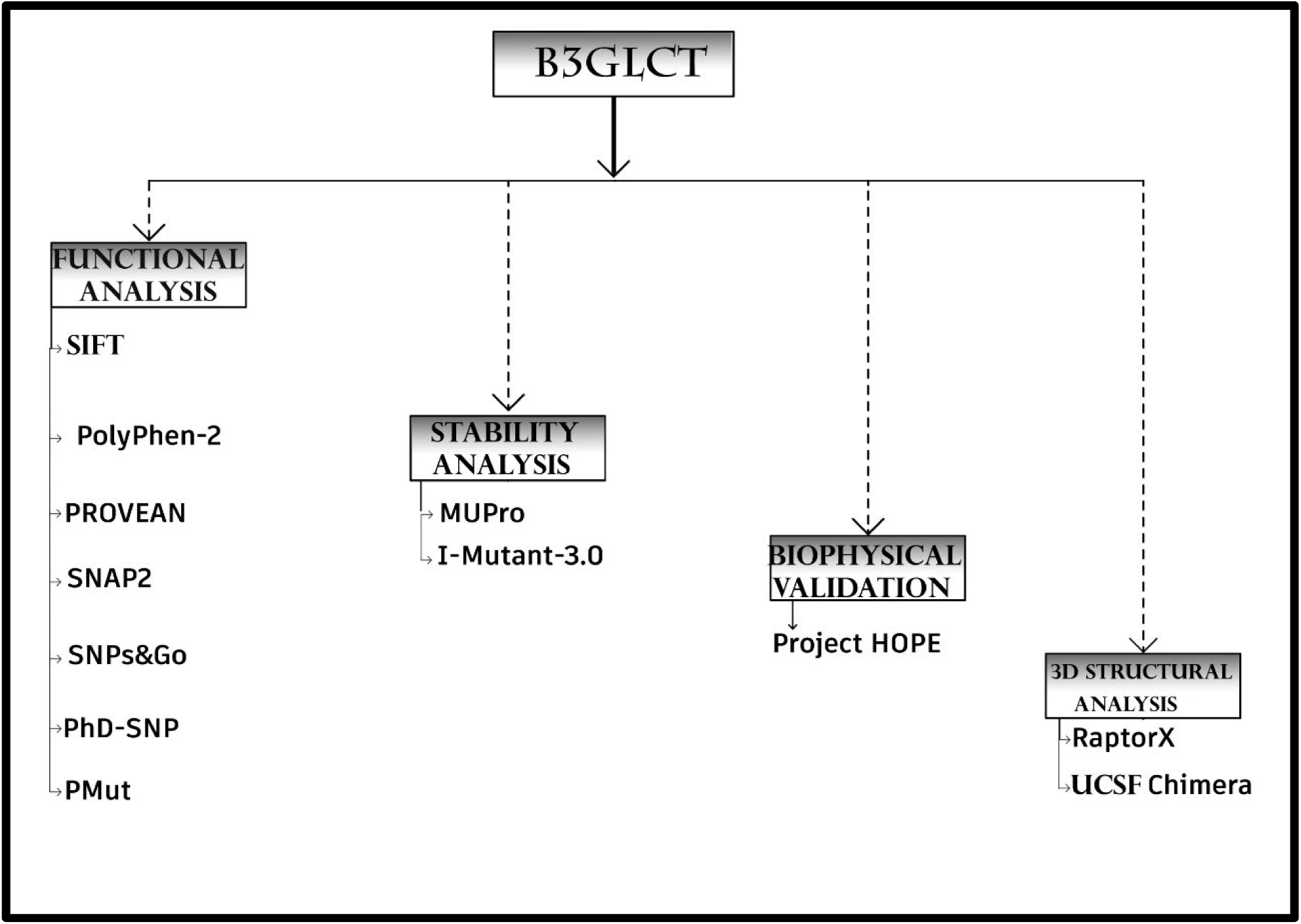
shows work flow of the paper (the type of analysis and soft wares used to analyses the SNPs in B3GLCT gene).

**Figure 2:**
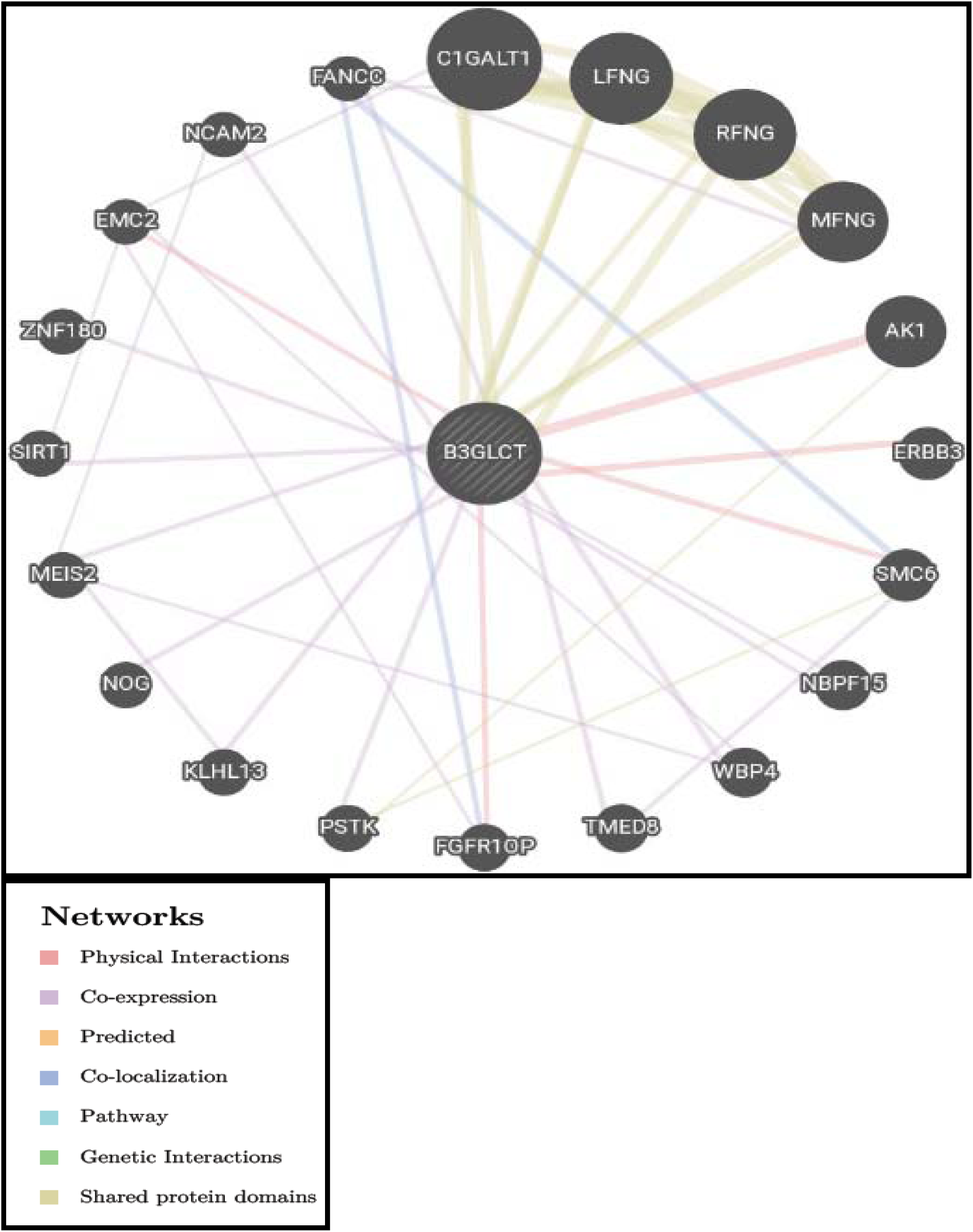
Interaction between B3GLCT and its related gene as predicted by gene MANIA.

## 4. Discussion

Thirteen novel single nucleotides polymorphism in B3GLCT gene has been discovered to be related to peters plus syndrome. Physiochemical and structural proprieties of these SNPs was analyzed and presented in this paper along with the predicted protein structural changes. (Figure 1)

Out of 1035 SNPs downloaded from NCBI, 371 SNPs were found to be missense mutations, we further analyzed these SNPS by SIFT,Polyphen2,Provean and SNAP2 which resulted in 40 SNPs(Table 1), which upon further analysis by SNPand GO and PMut resulted in 13 missense SNPs (Table 2). Imutant software was used to visualize the effect of these SNPs on the stability of the proteins, which shows six SNPs increasing the stability of the protein (Y172C, A222V, C260R, R377C, G379C, and G393E). While the other seven SNPs caused a decrease in the stability of the protein (C260Y, D349G, I354K, G393R, G395E, G425E and R445W). However all the 13 SNPs reduced the stability according to Mupro. This indicate the deleterious effect of these SNPs on the stability and hence the function of the protein.(42-44) (Table3).

**Table 3:** structural investigation calculated using I-Mutant3.0 and MUpro respectively.

| Mutations | SVM2 Prediction Effect | RI | DDG Value Prediction | Mupro Prediction | Mupro Score |
| --- | --- | --- | --- | --- | --- |
| Y172C | Increase | 3 | -0.75 | DECREASE | -0.82188586 |
| A222V | Increase | 2 | 0.23 | DECREASE | -0.76591056 |
| C260R | Increase | 0 | -0.23 | DECREASE | -1.3691514 |
| C260Y | Decrease | 0 | -0.14 | DECREASE | -1.2236219 |
| D349G | Decrease | 1 | -0.99 | DECREASE | -1.6758943 |
| I354K | Decrease | 6 | -1.97 | DECREASE | -1.7050794 |
| R377C | Increase | 1 | -0.73 | DECREASE | -0.91458682 |
| G379C | Increase | 1 | -0.79 | DECREASE | -0.63024675 |
| G393R | Decrease | 2 | -0.28 | DECREASE | -0.61407846 |
| G393E | Increase | 1 | -0.34 | DECREASE | -0.72438853 |
| G395E | Decrease | 1 | -0.65 | DECREASE | -0.350275 |
| G425E | Decrease | 1 | -0.57 | DECREASE | -0.59914297 |
| R445W | Decrease | 5 | -0.35 | DECREASE | -1.4166192 |

Physiochemical properties was analyzed using Hope project software, which also provided us with 2D structural prediction of wild and mutant amino acids. In addition, it predicted the following for Y172C: differences in size between the wild and mutant amino acids with the mutant amino acids being smaller witch may lead to loos of interaction especially when combined with the predicted increase in the mutant residue hydrophobicity. In case of A222V:

The mutant residue is predicted to be bigger than the wild one, which may lead to bumps.

Furthermore, we found in C260R SNP: an increase in the charges and size of the mutant amino acid with changes in the hydrophobicity, which disrupt interactions with other molecules. In C260Y: There is a difference in charge and hydrophobicity between the wild type and mutant amino acid, which results in repulsion of ligands or other residues with the same charge, and loss of hydrophobic interactions. Analysis of D349G reviled a loss of charges, decease in size and increase in the hydrophobicity of the mutant amino acids. Interactions in I354K is lost due to changes in the charge and hydrophobicity and increased size of the mutant amino acids. Same changes are noted in R377C with decrease rather than increase size in the mutant amino acids. In G379C: the increase in size of the mutant amino acids may lead to bumps. G393R has the same characteristic and in addition, the mutant amino acid introduce a charge that can cause repulsion of ligands or other residues with the same charge. Furthermore, in G393E, G395E, G425E and R445W similar changes to G393R are noted.

Hope project also reported most pf the SNPs to exit in following conserved domains (Interpro Domain, Fringe-Like IPR003378 and Nucleotide-Diphospho-Sugar Transferases IPR029044) which indicate the significance of the mutations for the function of the protein.(45) The following 3 SNPS (A222V, C260R and C260Y) has not been reported to share the same domains.

Chimera software was used to visualize the changes in the structure of the wild and mutant amino acids, we combined the designed 3D image of chimera with the generated 2D images from Hope project website which further proves the significant changes inflicted by these thirteen mutants.(Figure4- 16). Also the full predicted structure of the predicted was visualized by chimera (Figure 3).(supplementary file 1).

**Figure (3):**
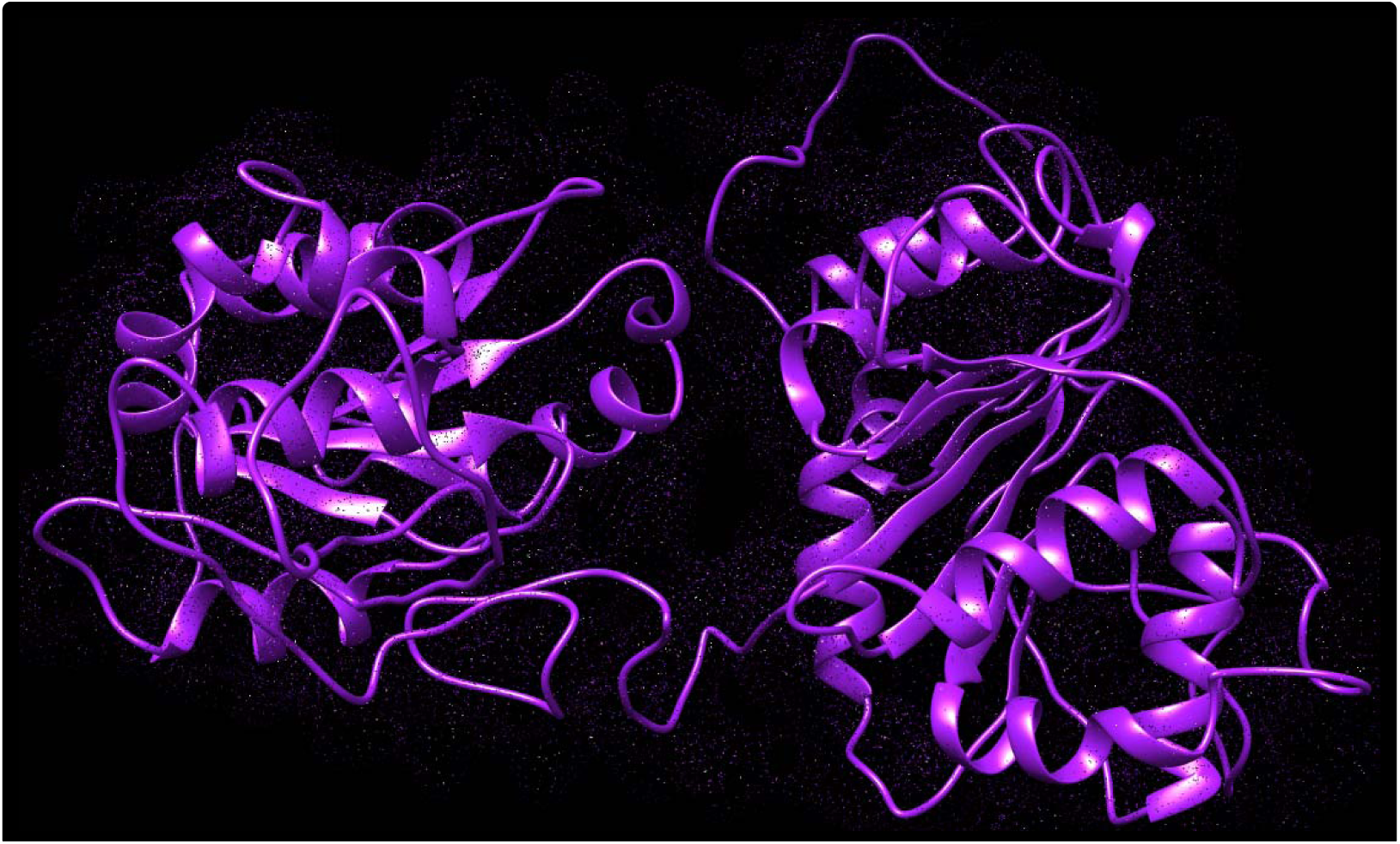
Full structure of B3GALTL gene related protein as predicted by Chimera software. Purple color indicates the background structure of the protein predicted by Chimera software.

**Figure (4):**
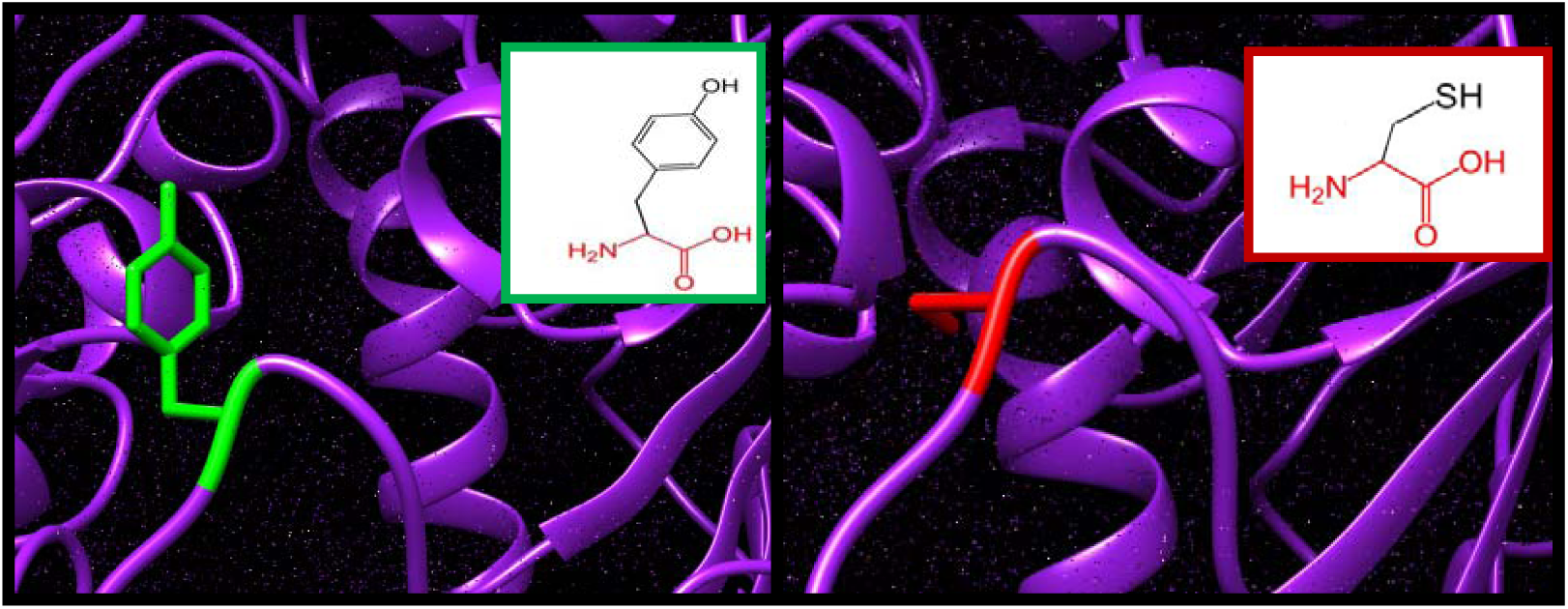
Y172C: Threonine amino acid changed to Cysteine at position 172 in B3GALTL gene related Protein as predicted by Chimera and Hope soft wares. Green color indicates wild amino acids and red color indicates mutant amino acid predicted by Chimera software. Green small box indicates 2D wild amino acid and red small box indicates 2D mutants amino acid predicted by Hope project software. Purple color indicates the background structure of the protein predicted by Chimera software.

**Figure (5):**
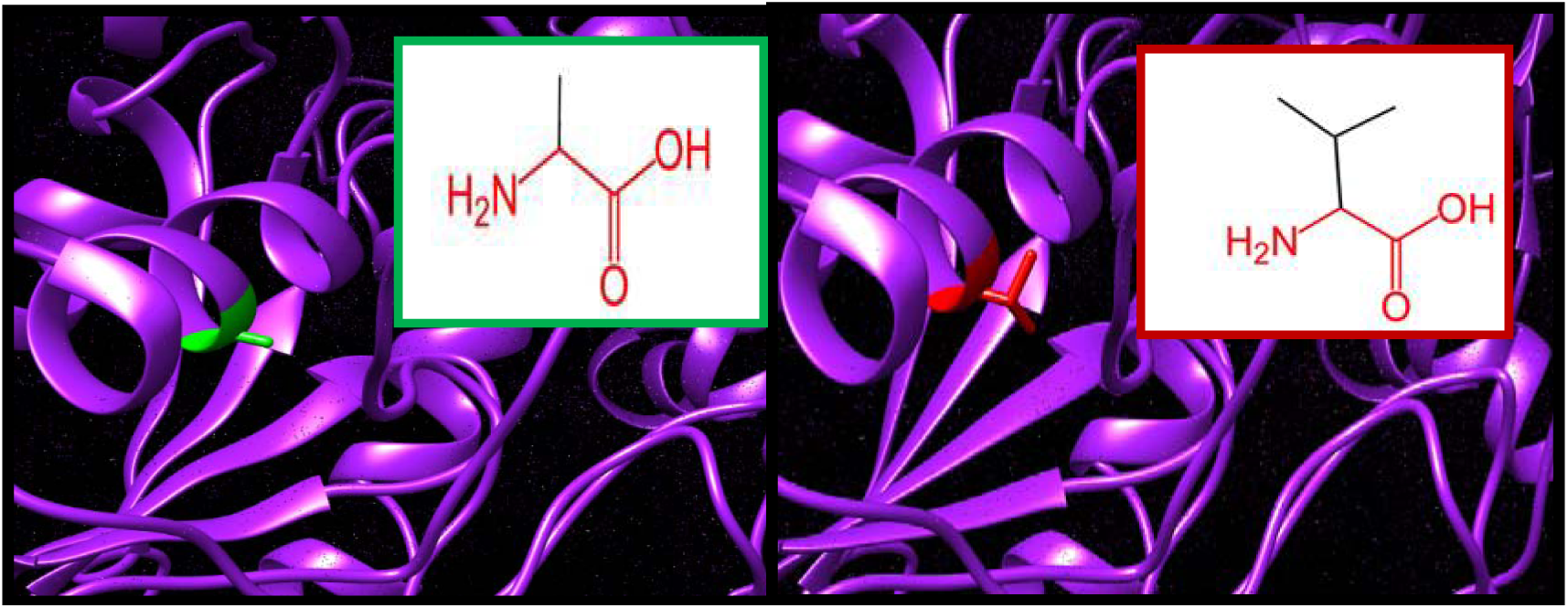
A222V: Alanine amino acid changed to Valine at position 222 in B3GALTL gene related Protein as predicted by Chimera and Hope soft wares. Green color indicates wild amino acids and red color indicates mutant amino acid predicted by Chimera software. Green small box indicates 2D wild amino acid and red small box indicates 2D mutants amino acid predicted by Hope project software. Purple color indicates the background structure of the protein predicted by Chimera software.

**Figure (6):**
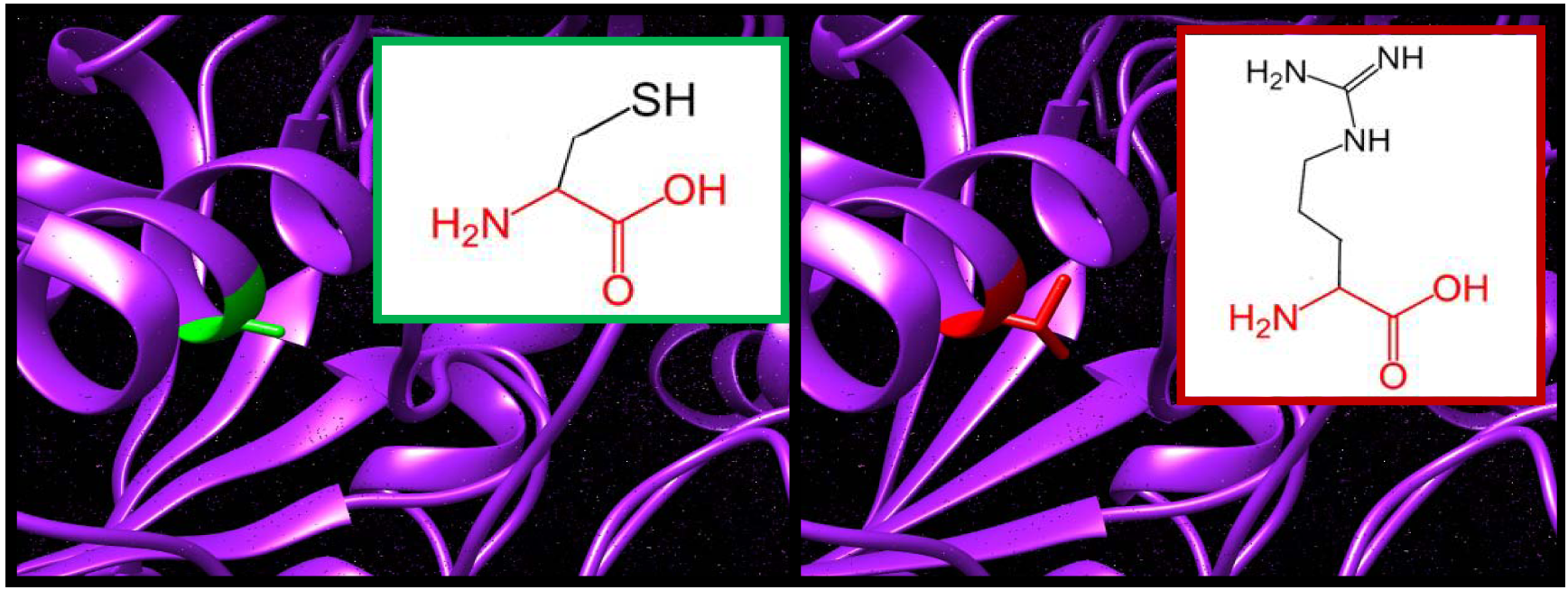
C260R: Cysteine amino acid changed to Arginine at position 260 in B3GALTL gene related Protein as predicted by chimera and Hope soft wares. Green color indicates wild amino acids and red color indicates mutant amino acid predicted by Chimera software. Green small box indicates 2D wild amino acid and red small box indicates 2D mutants amino acid predicted by Hope project software. Purple color indicates the background structure of the protein predicted by Chimera software.

**Figure (7):**
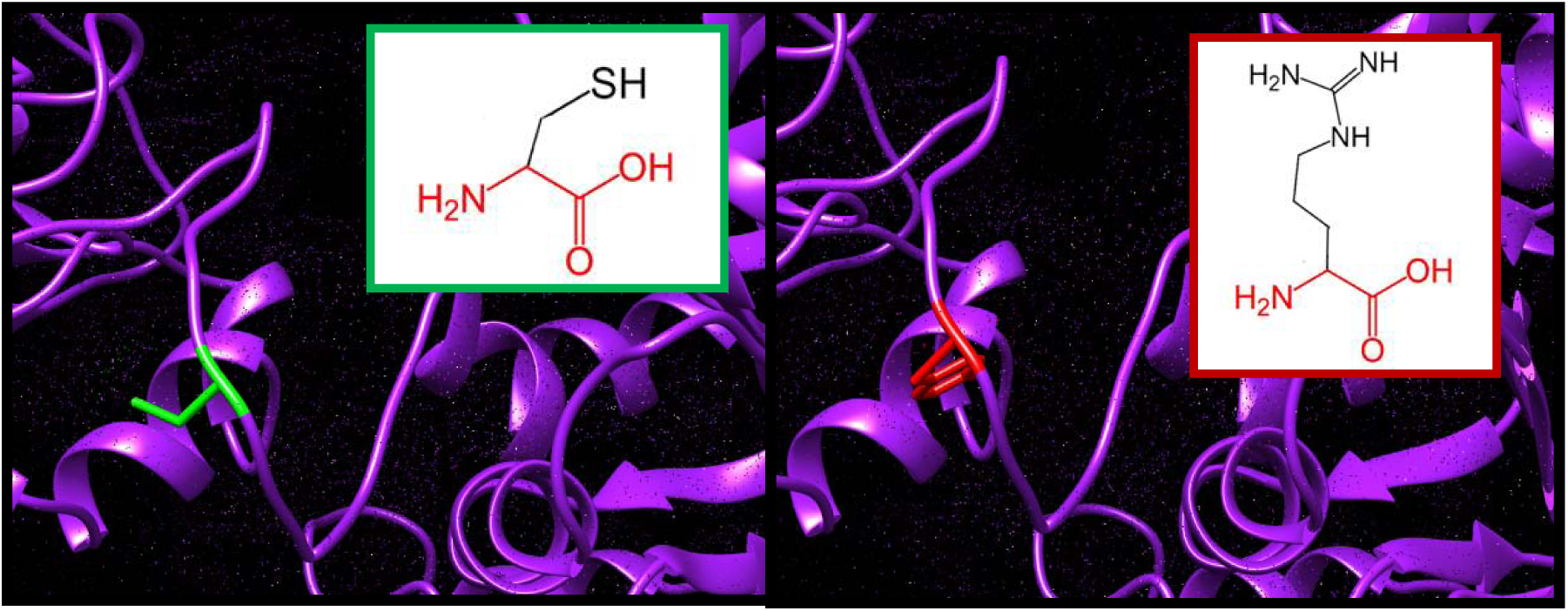
C260Y: Cysteine amino acid changed to Tyrosine in position 260 in B3GALTL gene related Protein as predicted by Chimera and Hope soft wares. Green color indicates wild amino acids and red color indicates mutant amino acid predicted by Chimera software. Green small box indicates 2D wild amino acid and red small box indicates 2D mutants amino acid predicted by Hope project software. Purple color indicates the background structure of the protein predicted by Chimera software.

**Figure (8):**
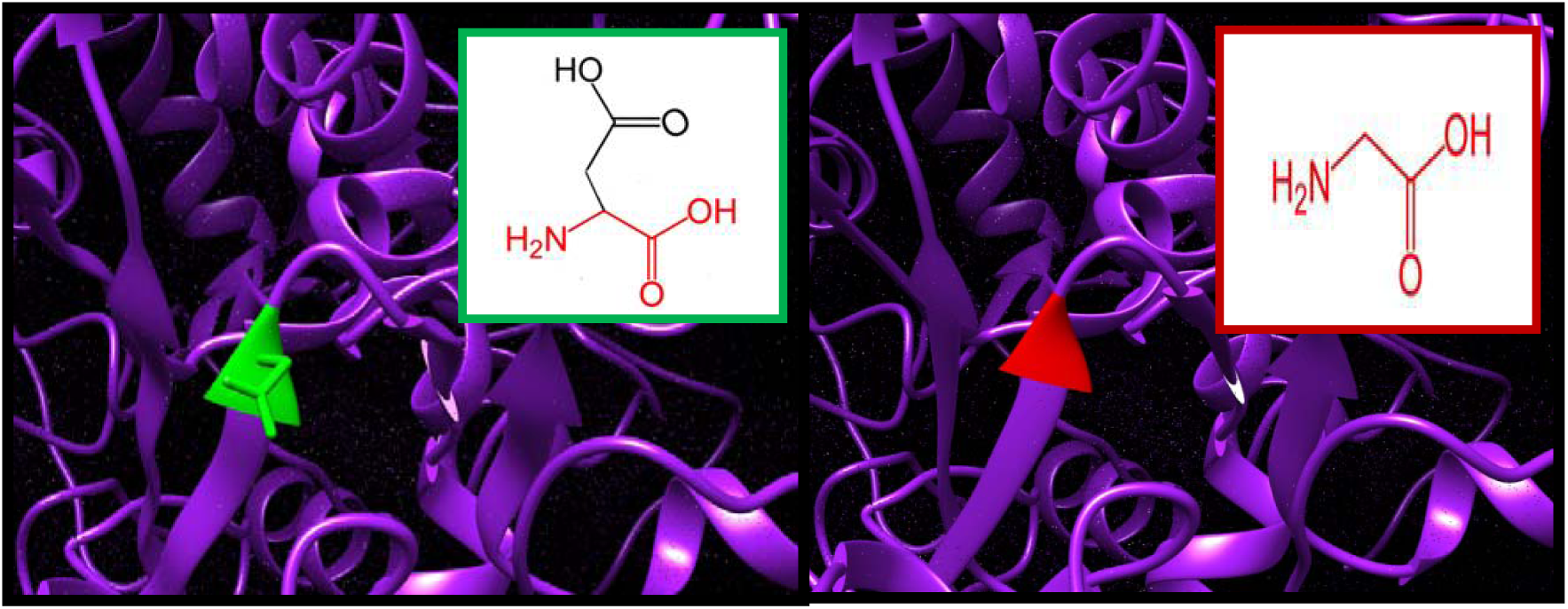
D349G: Aspartic acid amino acid changes to Glycine in position 349 in B3GALTL gene related Protein as predicted by Chimera and Hope soft wares. Green color indicates wild amino acids and red color indicates mutant amino acid predicted by Chimera software. Green small box indicates 2D wild amino acid and red small box indicates 2D mutants amino acid predicted by Hope project software. Purple color indicates the background structure of the protein predicted by Chimera software.

Gene MAINA was used to study the association between B3GLCT gene and other genes which shows that the gene co-expressed and physically interact with many genes like (NBPF15, FANCC, MEIS2 and TMED8) and (SMC6, FGFR1OP, EMC2 and ERBB3) respectively. (Table 4)

**Table (4):** The gene co-expressed, share domain and Interaction with B3GLCT gene network:

| Gene 1 | Gene 2 | Weight | Network group |
| --- | --- | --- | --- |
| **NBPF15** | **B3GLCT** | **0.009724106** | **Co-expression** |
| **EMC2** | **FGFR1OP** | **0.007030725** | **Co-expression** |
| **FANCC** | **B3GLCT** | **0.019513212** | **Co-expression** |
| **MEIS2** | **WBP4** | **0.008299122** | **Co-expression** |
| **TMED8** | **SMC6** | **0.017977167** | **Co-expression** |
| **NBPF15** | **B3GLCT** | **0.019043844** | **Co-expression** |
| **WBP4** | **B3GLCT** | **0.03277153** | **Co-expression** |
| **TMED8** | **B3GLCT** | **0.027544448** | **Co-expression** |
| **ZNF180** | **B3GLCT** | **0.023338769** | **Co-expression** |
| **FANCC** | **MFNG** | **0.021766916** | **Co-expression** |
| **PSTK** | **B3GLCT** | **0.026062034** | **Co-expression** |
| **SIRT1** | **B3GLCT** | **0.028160973** | **Co-expression** |
| **NCAM2** | **B3GLCT** | **0.02477317** | **Co-expression** |
| **KLHL13** | **B3GLCT** | **0.028947689** | **Co-expression** |
| **NOG** | **B3GLCT** | **0.027127726** | **Co-expression** |
| **MEIS2** | **B3GLCT** | **0.028704958** | **Co-expression** |
| **MEIS2** | **KLHL13** | **0.02369096** | **Co-expression** |
| **EMC2** | **C1GALT1** | **0.00674697** | **Co-expression** |
| **EMC2** | **WBP4** | **0.009009355** | **Co-expression** |
| **EMC2** | **SIRT1** | **0.004835495** | **Co-expression** |
| **NCAM2** | **MEIS2** | **0.005485816** | **Co-expression** |
| **FANCC** | **LFNG** | **0.009594724** | **Co-expression** |
| **FANCC** | **SMC6** | **0.025926804** | **Co-localization** |
| **FANCC** | **FGFR1OP** | **0.017828617** | **Co-localization** |
| **AK1** | **B3GLCT** | **0.17603359** | **Physical Interactions** |
| **SMC6** | **B3GLCT** | **0.08657284** | **Physical Interactions** |
| **FGFR1OP** | **B3GLCT** | **0.062340308** | **Physical Interactions** |
| **EMC2** | **B3GLCT** | **0.055467047** | **Physical Interactions** |
| **ERBB3** | **B3GLCT** | **0.2105723** | **Physical Interactions** |
| **C1GALT1** | **B3GLCT** | **0.110943414** | **Shared protein domains** |
| **LFNG** | **B3GLCT** | **0.077230476** | **Shared protein domains** |
| **LFNG** | **C1GALT1** | **0.30208087** | **Shared protein domains** |
| **RFNG** | **B3GLCT** | **0.077230476** | **Shared protein domains** |
| **RFNG** | **C1GALT1** | **0.30208087** | **Shared protein domains** |
| **RFNG** | **LFNG** | **0.4469795** | **Shared protein domains** |
| **MFNG** | **B3GLCT** | **0.039322563** | **Shared protein domains** |
| **MFNG** | **C1GALT1** | **0.107779875** | **Shared protein domains** |
| **MFNG** | **LFNG** | **0.15950671** | **Shared protein domains** |
| **MFNG** | **RFNG** | **0.15950671** | **Shared protein domains** |
| **PSTK** | **AK1** | **0.010328047** | **Shared protein domains** |
| **PSTK** | **SMC6** | **0.010580378** | **Shared protein domains** |
| **C1GALT1** | **B3GLCT** | **0.25** | **Shared protein domains** |
| **LFNG** | **B3GLCT** | **0.25** | **Shared protein domains** |
| **LFNG** | **C1GALT1** | **0.25** | **Shared protein domains** |
| **RFNG** | **B3GLCT** | **0.25** | **Shared protein domains** |
| **RFNG** | **C1GALT1** | **0.25** | **Shared protein domains** |
| **RFNG** | **LFNG** | **0.25** | **Shared protein domains** |
| **MFNG** | **B3GLCT** | **0.25** | **Shared protein domains** |
| **MFNG** | **C1GALT1** | **0.25** | **Shared protein domains** |
| **MFNG** | **LFNG** | **0.25** | **Shared protein domains** |
| **MFNG** | **RFNG** | **0.25** | **Shared protein domains** |

Previous researches reported c.660+1G>A(rs80338851) as the most common mutation in Peters plus syndrome (PPS)(12, 22). While other work done by Weh et al. reported different types of mutations, one of them is a missense mutation (G394E) which was also reported to have deleterious effect in our analysis by the first four soft wares (SIFT, Provean,Polyphen2 and SNAP2).

These thirteen SNPs play major role in the advancement of therapeutic and diagnostic techniques related to the Peters plus syndrome which hopefully will lead to better prognosis. However, this study is limited with the few number of available literature about the syndrome and the genes involved in it. Further wet lab studies are recommend to confirm the significance of these thirteen SNPs and to clarify its pathogenic role in the development of peters plus syndrome.

## Conclusion

The study of single nucleotide polymorphisms (SNPs) mutation in B3GLCT gene was investigated using several functional and stability in silico analysis tools. It was found that the most deleterious nonsense sing SNPs mutations are Y172C, A222V, C260R, C260Y, D349G, I354K, R377C, G379C, G393R, G393E,G395E, G425E and R445W and they can be used as a diagnostic marker for Peters plus syndrome.

