## supplemental figures for "*In Silico* Analysis of *B3GALTL* Gene Reveling 13 Novel Mutations Associated with Peters’-plus syndrome"

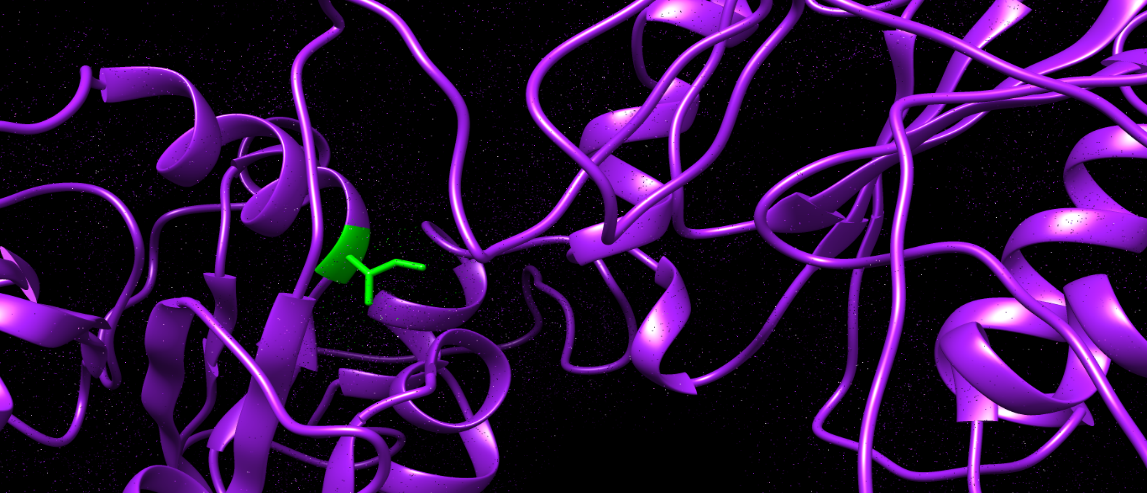

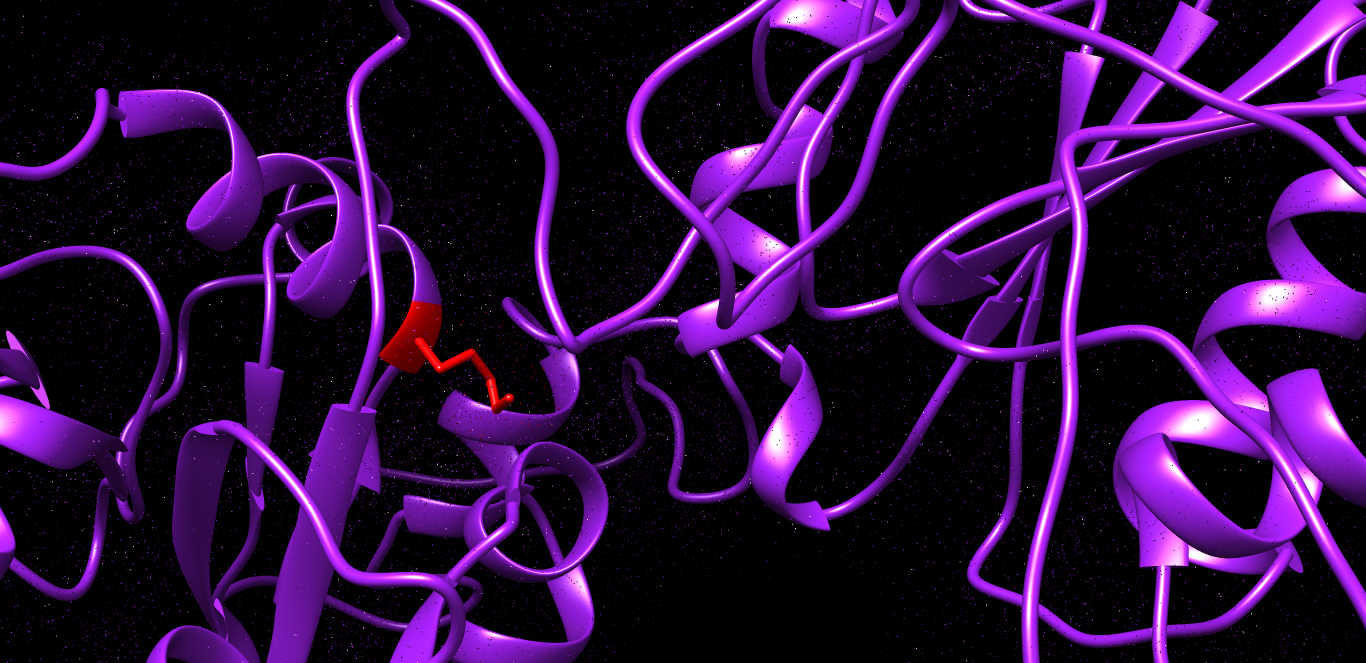

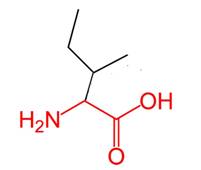

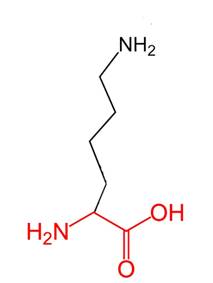

Figure (9): I354K: Isoleucine amino acid changed to Lysine in 354 in B3GALTL gene related Protein as predicted by chimera and Hope soft wares. Green color indicates wild amino acids and red color indicates mutant amino acid predicted by Chimera software. Green small box indicates 2D wild amino acid and red small box indicates 2D mutants amino acid predicted by Hope project software. Purple color indicates the background structure of the protein predicted by Chimera software.

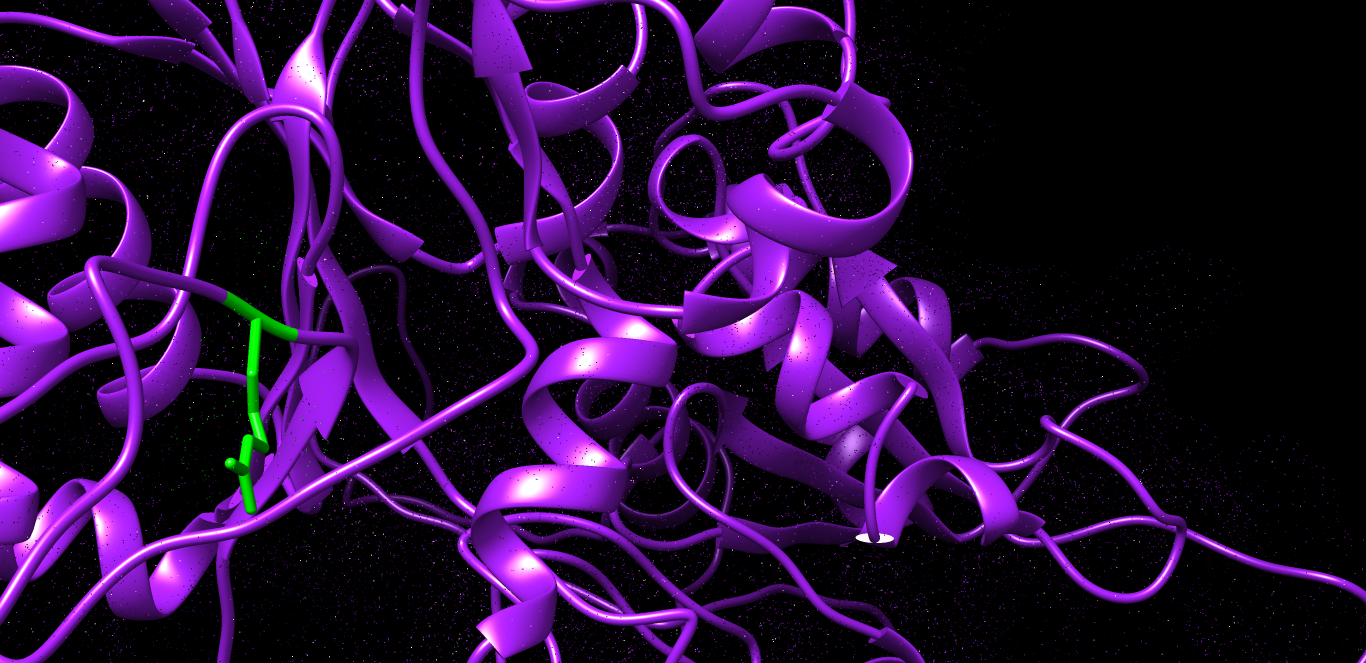

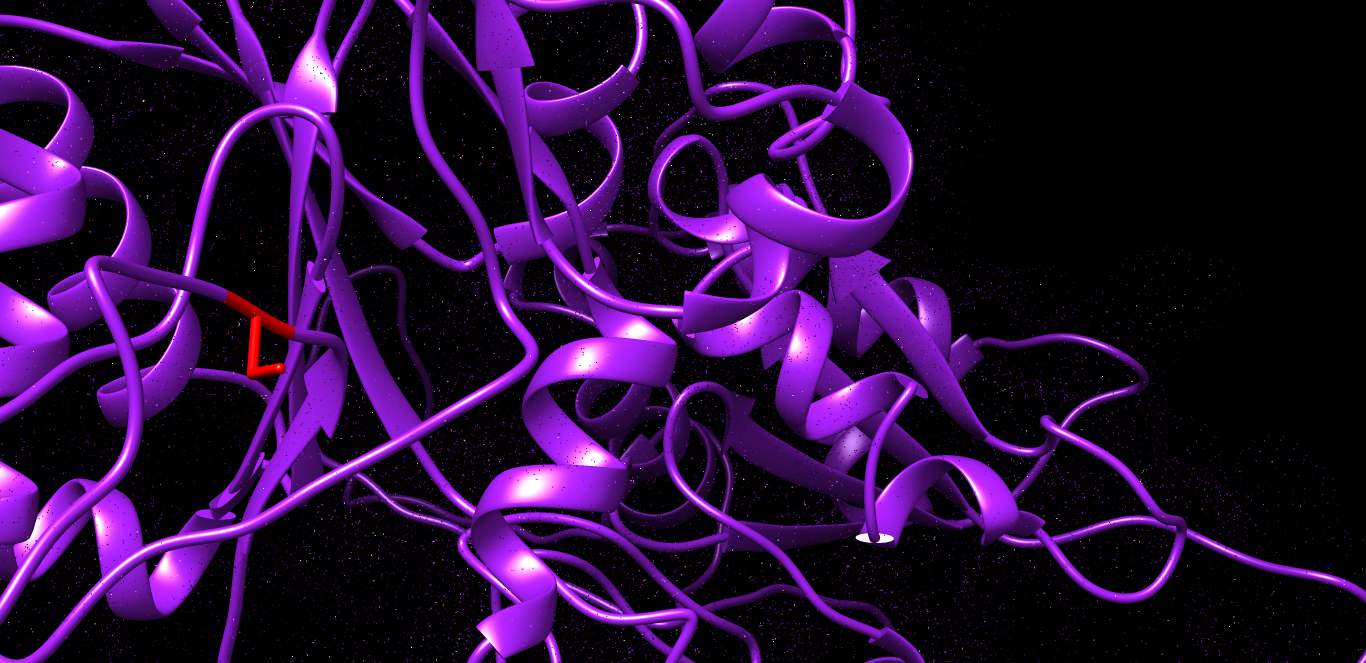

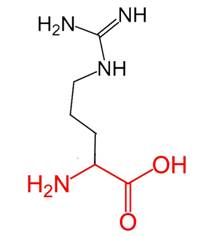

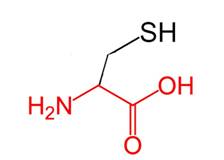

Figure (10): R377C: Arginine amino acid changed to Cysteine in position 377 in B3GALTL gene related Protein as predicted by Chimera and Hope soft wares. Green color indicates wild amino acids and red color indicates mutant amino acid predicted by Chimera software. Green small box indicates 2D wild amino acid and red small box indicates 2D mutants amino acid predicted by Hope project software. Purple color indicates the background structure of the protein predicted by Chimera software.

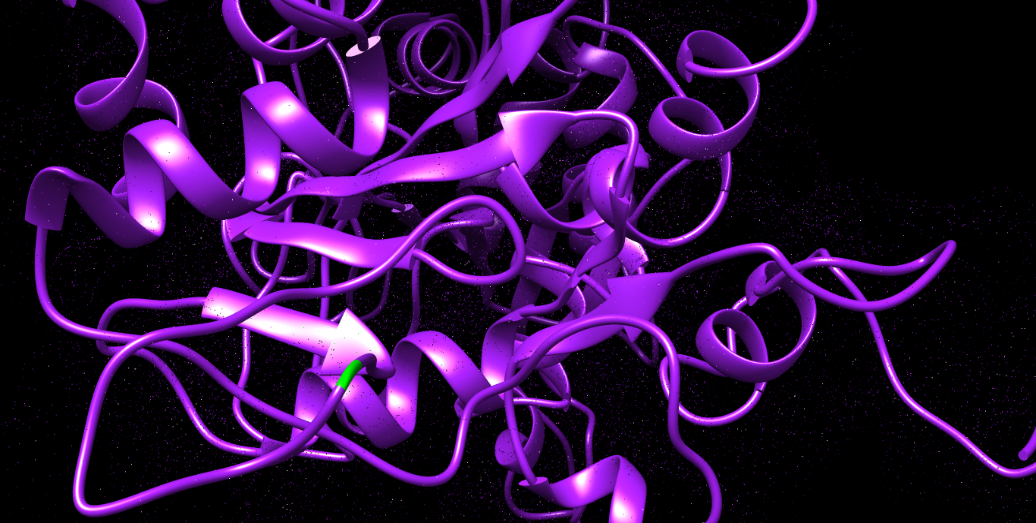

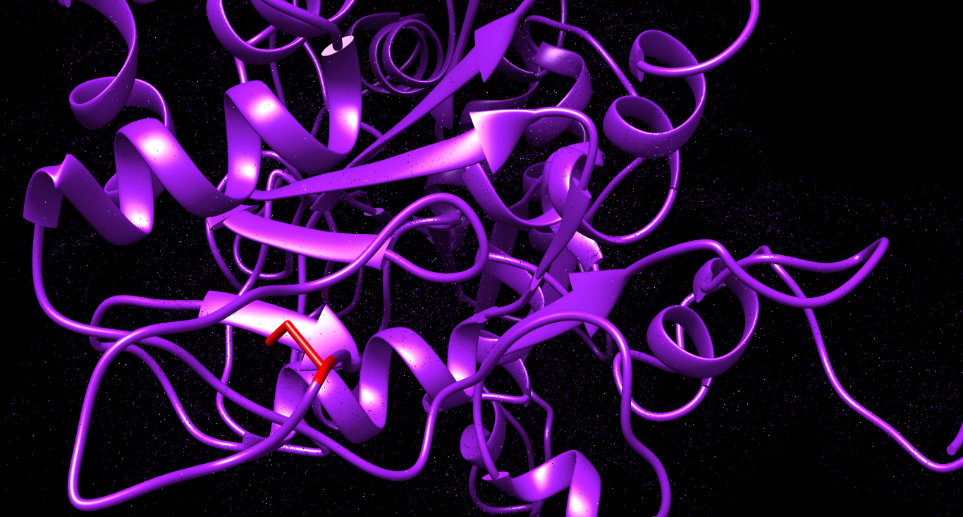

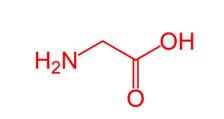

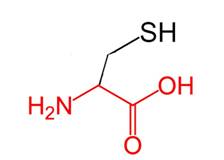

Figure (11): G379C: Glycine amino acid changed to Cysteine at position 379 in B3GALTL gene related Protein as predicted by Chimera and Hope soft wares. Green color indicates wild amino acids and red color indicates mutant amino acid predicted by Chimera software. Green small box indicates 2D wild amino acid and red small box indicates 2D mutants amino acid predicted by Hope project software. Purple color indicates the background structure of the protein predicted by Chimera software.

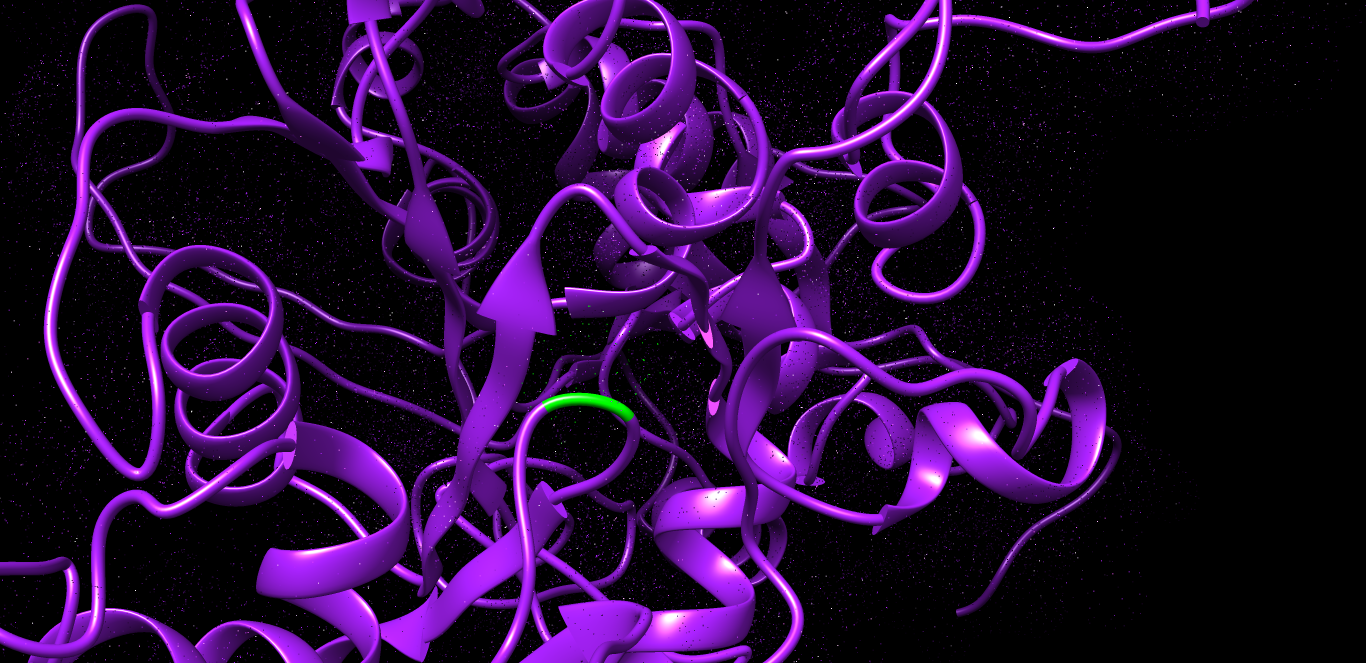

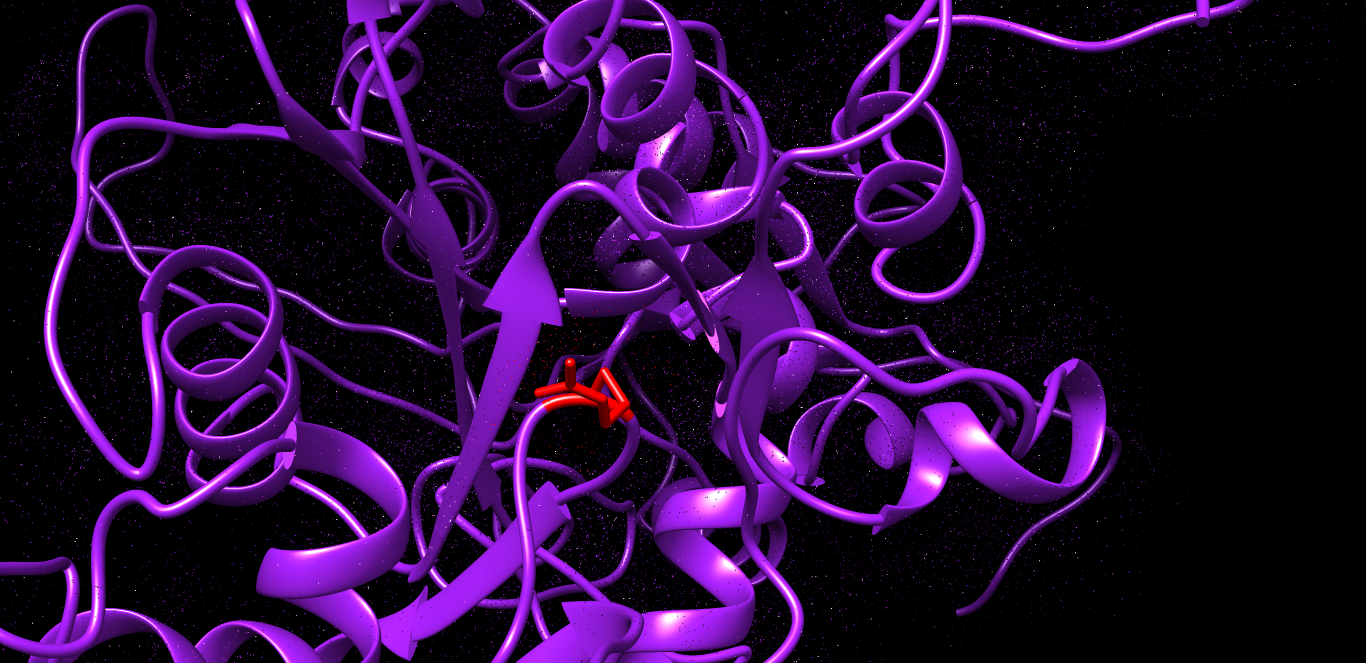

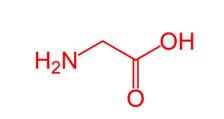

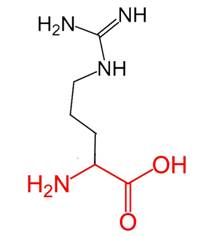

Figure (12): G393R: Glycine amino acid changed to Arginine in position 393 in B3GALTL gene related Protein as predicted by Chimera and Hope soft wares. Green color indicates wild amino acids and red color indicates mutant amino acid predicted by Chimera software. Green small box indicates 2D wild amino acid and red small box indicates 2D mutants amino acid predicted by Hope project software. Purple color indicates the background structure of the protein predicted by Chimera software.

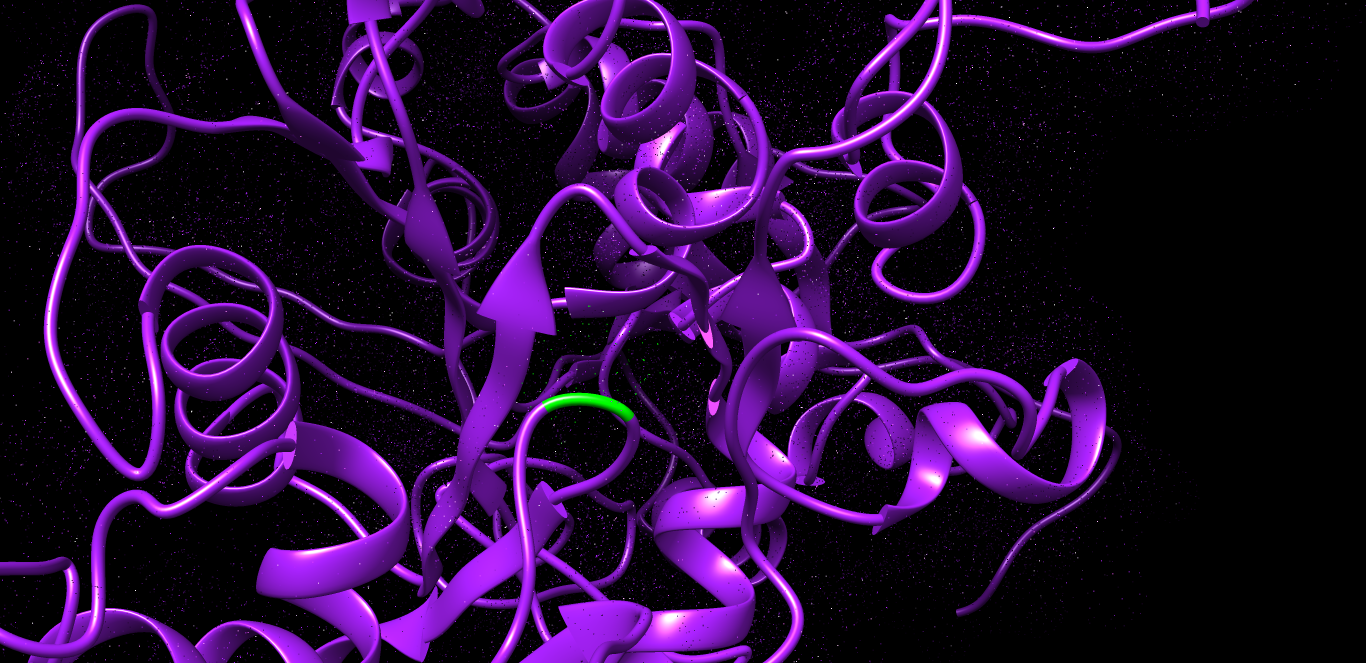

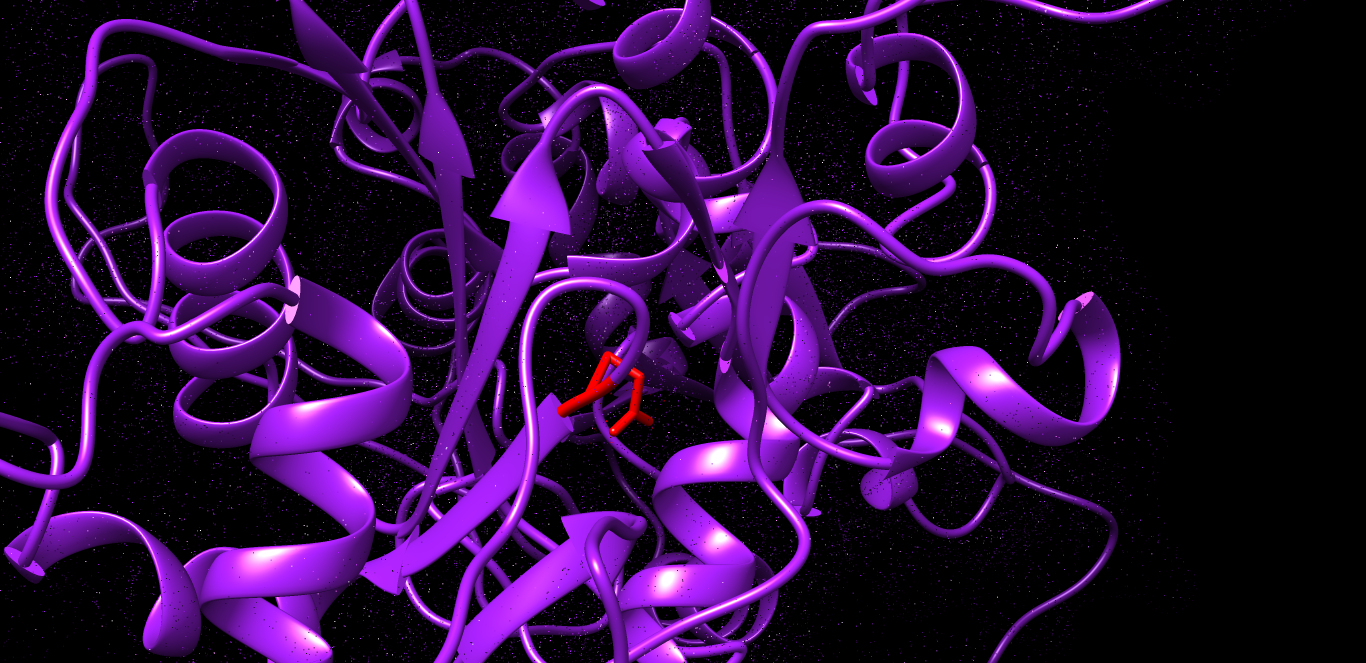

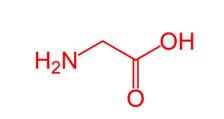

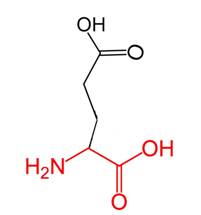

Figure (13): G393E: Glycine amino acid changed to Glutamic acid in position 393 in B3GALTL gene related Protein as predicted by Chimera and Hope soft wares. Green color indicates wild amino acids and red color indicates mutant amino acid predicted by Chimera software. Green small box indicates 2D wild amino acid and red small box indicates 2D mutants amino acid predicted by Hope project software. Purple color indicates the background structure of the protein predicted by Chimera software.

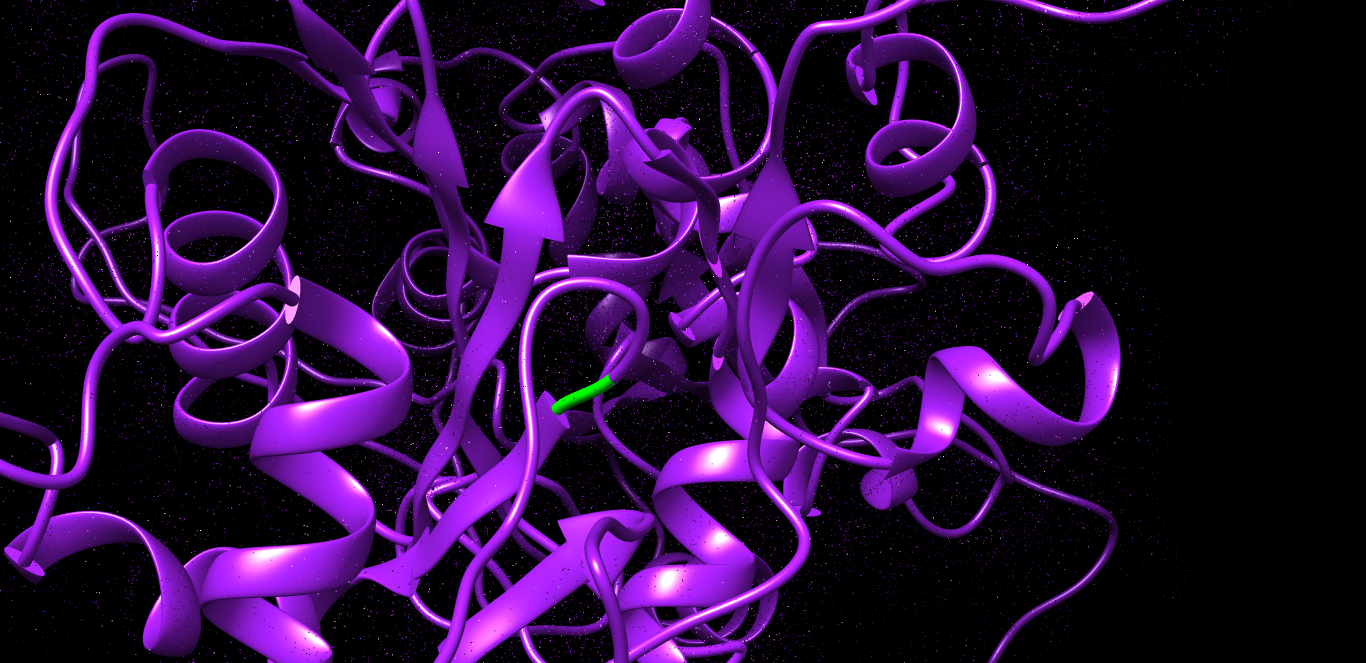

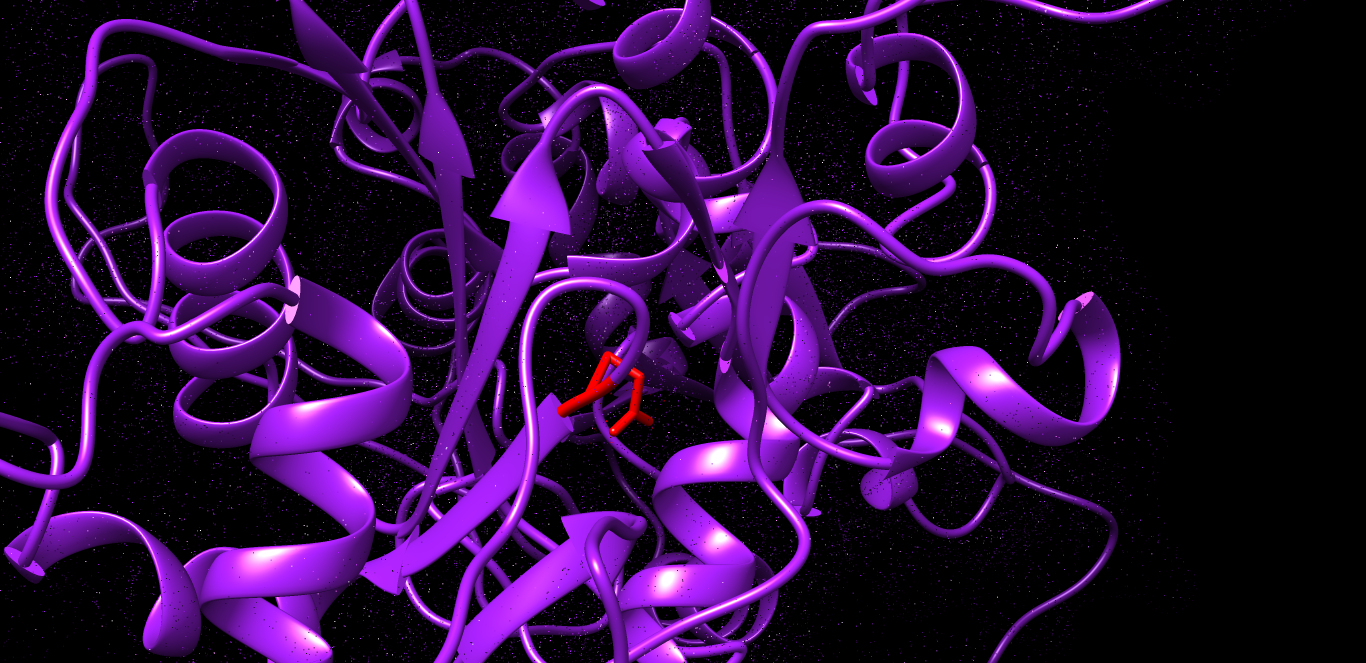

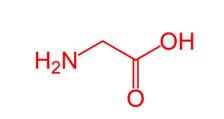

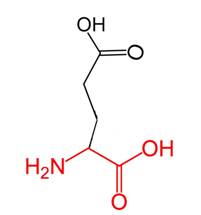

Figure (14): G395E: Glycine amino acid changed to Glutamic acid in position 359 in B3GALTL gene related Protein as predicted by Chimera and Hope soft wares. Green color indicates wild amino acids and red color indicates mutant amino acid predicted by Chimera software. Green small box indicates 2D wild amino acid and red small box indicates 2D mutants amino acid predicted by Hope project software. Purple color indicates the background structure of the protein predicted by Chimera software.

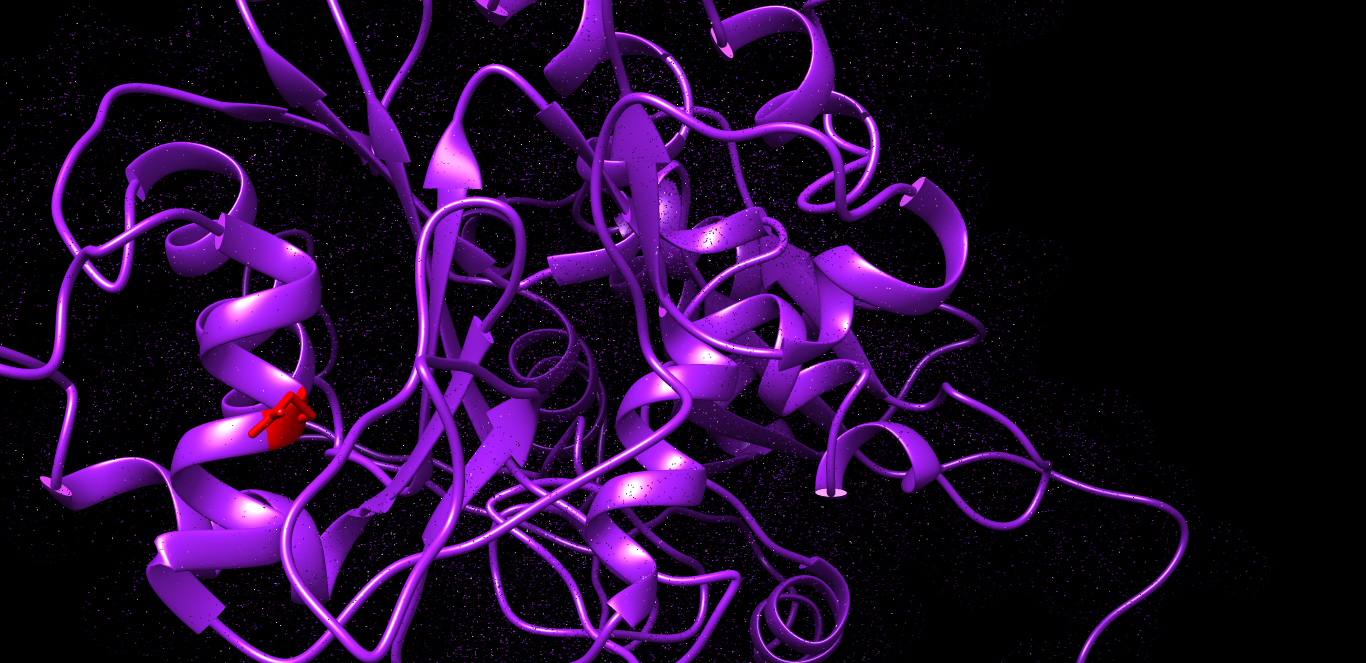

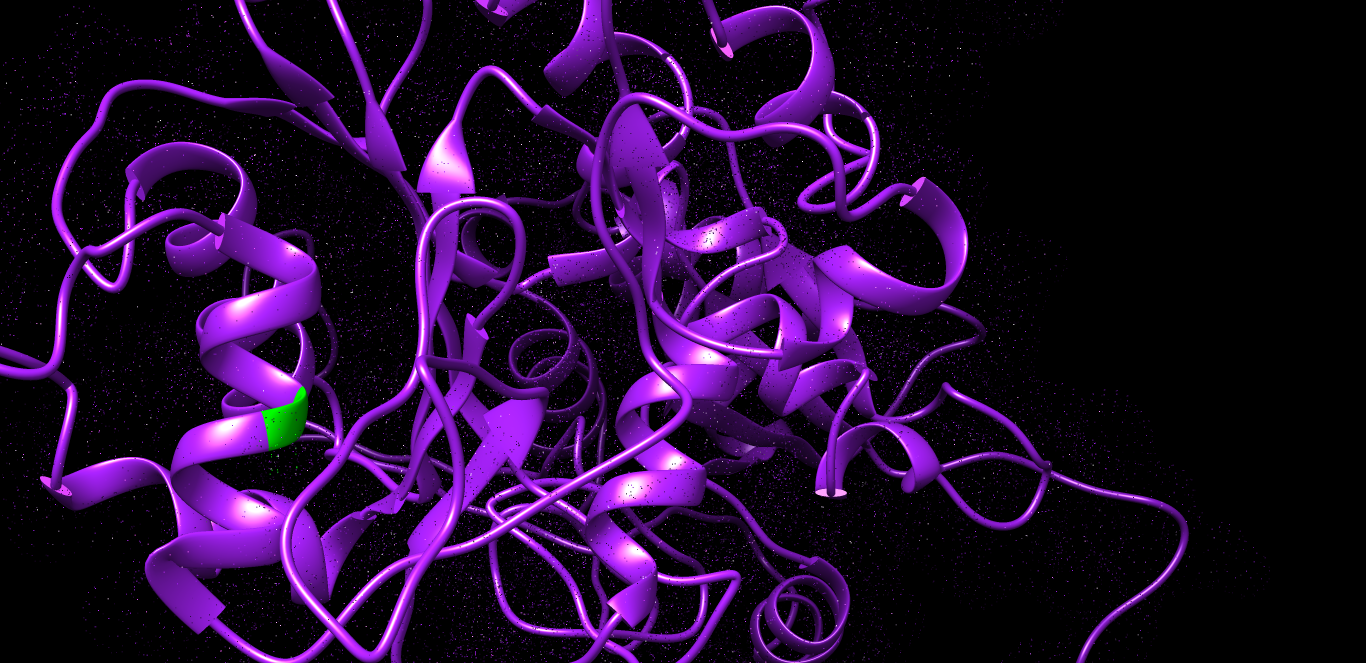

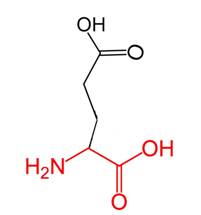

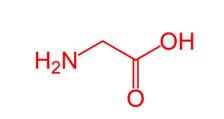

Figure (15): G425E: Glycine amino acid changed to Glutamic acid in position 425 in B3GALTL gene related Protein as predicted by Chimera and Hope soft wares. Green color indicates wild amino acids and red color indicates mutant amino acid predicted by Chimera software. Green small box indicates 2D wild amino acid and red small box indicates 2D mutants amino acid predicted by Hope project software. Purple color indicates the background structure of the protein predicted by Chimera software.

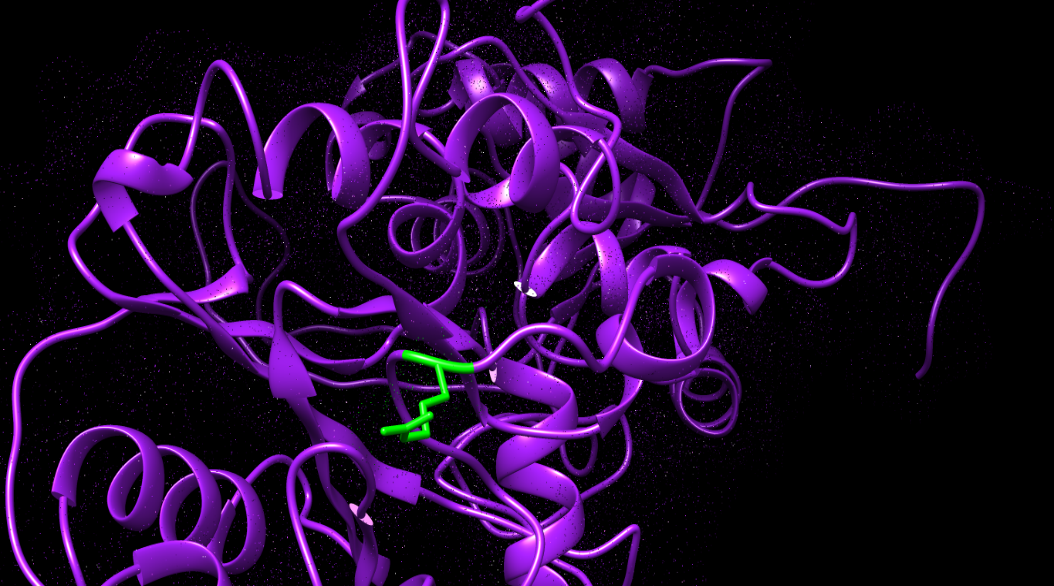

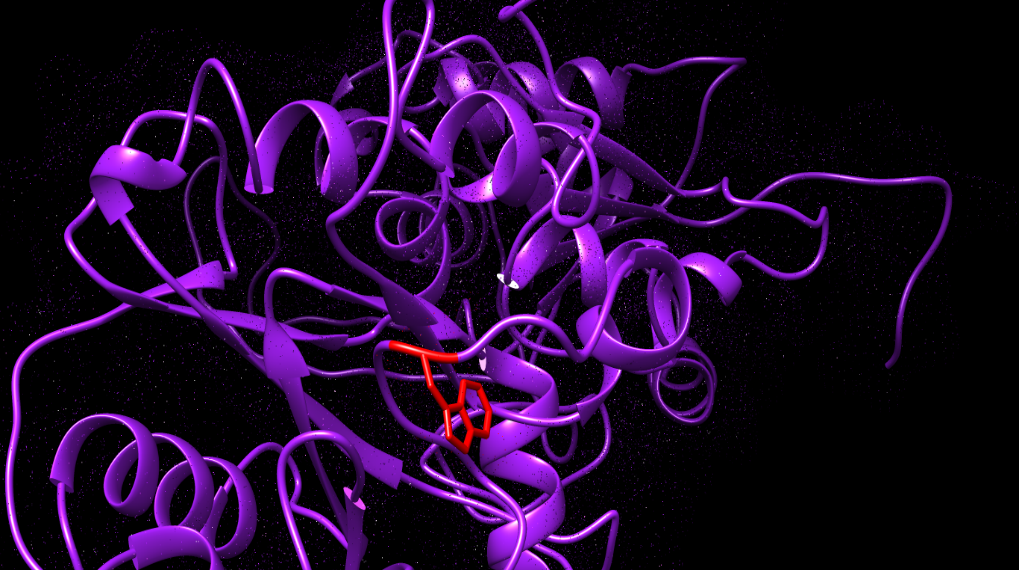

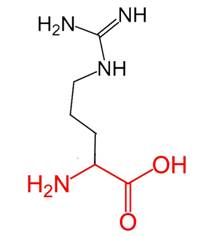

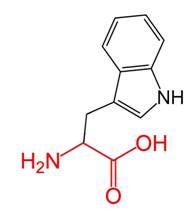

Figure (16): R445W: Arginine amino acid changed to Tryptophan in position 445 in B3GALTL gene related Protein as predicted by Chimera and Hope soft wares. Green color indicates wild amino acids and red color indicates mutant amino acid predicted by Chimera software. Green small box indicates 2D wild amino acid and red small box indicates 2D mutants amino acid predicted by Hope project software. Purple color indicates the background structure of the protein predicted by Chimera software
